# Drought tolerance is associated with constitutive gene expression, not plasticity, across California oak species

**DOI:** 10.1101/2025.08.19.671120

**Authors:** Alayna Mead, Camila D. Medeiros, Marissa E. Ochoa, Lawren Sack, Victoria L. Sork

## Abstract

- Drought is a major stressor for plants globally. Variation in gene expression patterns across species can provide critical evidence for the genomic basis of drought tolerance.
- We paired comparative transcriptomics with functional trait measurements to identify genomic mechanisms associated with drought tolerance across six species from three oak clades in California, including a pair of species within each clade representing relatively mesic or xeric environments. We tested how plastic and constitutive gene expression patterns varied among species with contrasting drought tolerance traits. We also tested whether gene expression responses were decoupled from phylogenetic history, suggesting they have evolved multiple times as adaptations to species climate niches.
- Species with drought-tolerant traits exhibited lower levels of gene expression plasticity during leaf dehydration than drought-sensitive species, but showed signatures of positive selection on constitutive gene expression. Drought-sensitive species across clades converged in their patterns of plastic gene expression during dehydration, diverging from their more closely related drought tolerant species, suggesting that repeated evolution has shaped plastic gene expression responses to drought.
- Drought-tolerant oak species have evolved constitutive gene expression alongside drought tolerant functional traits, while drought-sensitive oak species have evolved similar plastic gene expression responses to drought.

## Introduction

Drought is a major stressor for plants, and variation in aridity across environments can drive species distributions (Medeiros *et al*., 2023) and evolutionary diversification (Stebbins, 1952; Hipp *et al*., 2018). Climate change is increasing drought severity in some regions, including California (Diffenbaugh *et al*., 2015; IPCC, 2023), threatening plant populations that are not well-adapted to these new conditions and causing changes in ecosystem composition (Dong *et al*., 2019; Fettig *et al*., 2019; Taylor *et al*., 2020; Lemmo *et al*., 2024). Characterizing the underlying genetic basis of variation in drought responses among species can tell us how drought tolerance has evolved historically and how rapidly it might be able to evolve in existing populations. Understanding the evolution of drought tolerance requires studies linking traits and genomic data across taxa with varying tolerance. We investigated the genomic basis of drought tolerance in California oaks, a keystone lineage, using gene expression and phenotypic data. Specifically, we tested the hypothesis that species with similar levels of drought tolerance would exhibit similarities in their level of plasticity of gene expression, the genes and types of genes expressed, and their constitutive levels of gene expression.

The response of gene expression to leaf dehydration can be two-fold. As leaves dehydrate, genes should be expressed with a protective function preventing tissue damage, and also as a direct response to damage (e. g., to replace damaged proteins) (Chaves *et al*., 2003; Teskey *et al*., 2015; Moran *et al*., 2017). Drought tolerant species may show fewer changes in gene expression in response to drought because they are better able to withstand damage induced by leaf-level dehydration. For example, functional traits such as lower leaf wilting point, higher leaf mass per area, higher wood density, and higher water use efficiency can enable the maintenance of function during moderate droughts (Skelton *et al*., 2015; Anderegg *et al*., 2019). Alternatively, species with greater leaf-level drought tolerance may show greater expression of genes as part of an adaptive plastic response (Rivera *et al*., 2021). Comparing gene expression plasticity and drought tolerance traits among species can test whether drought tolerance at the leaf scale is broadly achieved through constitutive drought tolerance, such as that achieved by functional traits, or through rapid plasticity in gene expression during leaf dehydration.

Traditionally, comparisons of ecophysiological and functional traits among species have been used to understand where and when stress tolerance has evolved across a phylogeny (Kaproth & Cavender-Bares, 2016; Fletcher *et al*., 2018; Medeiros *et al*., 2023). However, such studies are limited by the number of traits that can be measured, and they typically represent average values for a species rather than plasticity over time (Swenson *et al*., 2017). Thus, measurements of traits alone may not represent species differences in response to stressors, nor provide insight into their genetic underpinnings (Ahrens *et al*., 2022). The use of gene expression responses to estimate drought tolerance could potentially expand the ability to estimate species drought tolerance within a lineage, either confirming that assessed using traits and current distributions, or indicating additional axes of drought tolerance. We thus tested whether species native to more arid habitats and with more drought tolerant traits show greater or lesser gene expression plasticity during leaf dehydration and differ in the types of genes being expressed. Evolutionary changes in gene expression are likely to be an important mechanism of adaptation (King & Wilson, 1975; Whitehead & Crawford, 2006; Gompel & Prud’homme, 2009; Jones *et al*., 2012; Stern, 2013). The amount of mRNA produced in a tissue under certain conditions can be considered a phenotype (Khaitovich *et al*., 2006) approximately corresponding to the amount of protein translated (Schwanhäusser *et al*., 2011). In contrast to traditional phenotypes, the measurement of gene expression makes it possible to identify specific genes, elucidating the genetic basis of drought-response traits. Here we focus specifically on genes that alter their expression under leaf drying, as they may be important in the plant’s plastic response to drought.

Drought tolerance may evolve through functional traits expressed in tissues that develop prior to the stress, and through adaptive phenotypic plasticity that occurs when the stress is experienced, but the relative importance of these mechanisms is unclear. Stress tolerance may evolve partly through changes in baseline levels of gene expression rather than plastic changes in the gene expression response to stress (Barshis *et al*., 2013; Akman *et al*., 2021; Collins *et al*., 2021; Le Provost *et al*., 2022; Dilworth *et al*., 2024). For example, frontloaded genes, which are continuously highly expressed, may confer stress tolerance in some species, while only being expressed in response to stress in other species (Rivera *et al*., 2021). Such constitutive expression could be evolutionarily advantageous if phenotypic plasticity has costs, such as maintaining sensory mechanisms, or limitations, such as the lag time between the environmental signal and the organismal response (DeWitt *et al*., 1998; Li *et al*., 2023). For example, constitutive expression of stress responses could be adaptive in environments that more frequently experience stressful conditions by eliminating the response lag time; or they could underlie development of morphological adaptations that have limited capacity for plasticity (Akman *et al*., 2021; Eisenring *et al*., 2022; Li *et al*., 2023). We tested whether variation in overall gene expression levels could be associated with drought tolerance traits by identifying genes that showed signatures of positive selection among species, and that were also correlated with drought-tolerance traits.

Drought tolerance across congeneric species are likely shaped by by parallel evolution of similar genes underlying the same phenotypes, but it is also possible that different genomic mechanisms result in the same phenotype through convergent evolution. If evolution tends to occur repeatedly using the same genes each time, then adaptation may be more constrained and occur slowly, but if some gene functions are redundant so that mutations in different genes can achieve the same phenotype, taxa may be more flexible in adapting to new conditions (Barrett & Schluter, 2008; Losos, 2011; Yeaman *et al*., 2016). Because definitions of parallel and convergent evolution differ across studies (Arendt & Reznick, 2008), we will use “parallelism” for cases when the same gene underlies an adaptive phenotype across species, “convergence” when different genes are used, and “repeated evolution” when the mechanism is unclear. Parallelism in gene expression may be common because regulatory regions can be evolutionary hotspots (Martin & Orgogozo, 2013) and is more likely to occur when only a few genes underlie an adaptive phenotype, simply because there are limited mechanisms allowing convergence to take place (O’Quin *et al*., 2010). However, in many cases, whole-organism level characteristics are determined by multiple genes, allowing a similar phenotype to evolve through multiple mechanisms (Savolainen *et al*., 2013; Yeaman *et al*., 2016; Agrawal, 2017; Lind *et al*., 2018). Because drought tolerance is thought to be polygenic, we predicted that responses to leaf drying would occur primarily through convergent changes at higher levels of biological organization than individual genes (Tenaillon *et al*., 2012; Renaut *et al*., 2014; Pfenninger *et al*., 2015; Bailey *et al*., 2015). Here, we quantify parallelism as the frequency at which ecologically similar but evolutionarily diverged species pairs responded to a treatment by altering expression of the same genes and test whether their responses to drying are more similar in the types of genes expressed than the individual genes.

Oaks (genus *Quercus*) are a diverse group of trees and shrubs that have adapted to a wide range of climates throughout their evolutionary history (Cavender-Bares *et al*., 2018; Kremer & Hipp, 2020; Sork *et al*., 2022). As the genus spread south from its paleo-arctic distribution over the last 50 million years, species have evolved traits that are beneficial for their new climate niches. Among those traits, drought tolerance is critical in niche adaptation of oak species (Abrams, 1990; Kaproth & Cavender-Bares, 2016; Ramírez-Valiente *et al*., 2018; Skelton *et al*., 2018; Lobo *et al*., 2018). Many North American oak species in different taxonomic sections of the genus have filled similar trait and climatic niches, and because species from different sections do not hybridize with each other (Nixon, 2002), we assume these traits arose independently within each section, or that species in multiple sections experienced similar selection pressures after divergence from a common ancestor. Distantly related oak species often co-occur in the same community, and these co-occurring species tend to have similar traits that they may have independently evolved while adapting to the same climate (Mohler, 1990; Cavender-Bares *et al*., 2004; Cavender-Bares *et al*., 2018). Additionally, phylogenetic data show that the two largest sections in North America, the red oaks (section *Lobatae*) and white oaks (section *Quercus*) have radiated into new climatic niches in parallel (Hipp *et al*., 2018), while species within *Protobalanus*, a small section that is geographically restricted to western North America, also inhabit different climate niches (Nixon, 2002).

The overarching goals of this study are to assess the extent to which gene expression responses to leaf dehydration vary across oaks in association with functional traits that confer drought tolerance, and further, to test whether gene expression reflects parallel or convergent evolution. We compared drought-related traits and gene expression responses to leaf drying across six oak species from three sections (clades within the genus *Quercus*), selecting two species from contrasting mesic or xeric climates within each section (Figure 1). We tested three hypotheses: H1. Plastic gene expression in response to dehydration is lower in species with drought tolerant traits and that are adapted to more arid climates. H2. Drought-tolerant species have evolved constitutive regulation of gene expression rather than an adaptive plastic response to dehydration. H3. Gene expression responses are decoupled from phylogenetic history and evolved multiple times as adaptations to species climate niches. We present the under-explored evidence that drought tolerance is associated with constitutive gene expression, while drought-sensitive species respond to drought through gene expression plasticity, and that for both strategies, some genes have evolved expression patterns that are independent of phylogenetic history.

**Figure 1.**
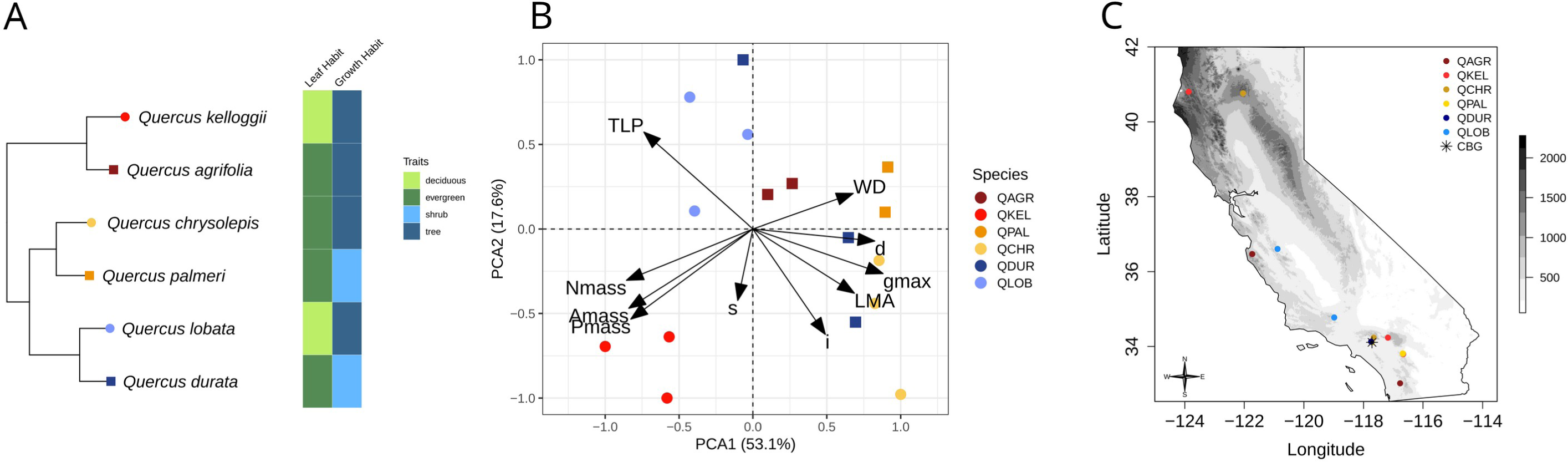
(A) Phylogeny of species, including their leaf habit and growth habit. Within each section, the more drought-tolerant species is indicated with a square, and the more drought-sensitive species with a circle. (B) PCA of mechanistic drought traits: leaf mass per area (LMA), wood density (WD), turgor loss point (TLP), leaf nitrogen and phosphorus content per mass (N_mass_, P_mass_), time-integrated photosynthetic rate per mass (A_mass_), stomatal density (d), stomatal initiation rate (i), and anatomical maximum stomatal conductance (g_max_). (C) Location of provenance origins for each species and location of the common garden (California Botanic Garden, CBG).

## Materials and Methods

### Study species

We sampled six of the 21 species of California native oaks, with two representatives from each of the three *Quercus* sections found in North America, *Quercus*, *Lobatae*, and *Protobalanus* (Hipp *et al*., 2018). Within each of the three sections, we chose two representative species – a relatively drought-sensitive and drought-tolerant species (Figure 1A). Within section *Quercus*, the white oaks, *Q. lobata* is a large deciduous tree species occupying grasslands and riparian habitats and is expected to be drought-sensitive; while *Q. durata* var. *gabrielensis* is an evergreen scrub oak occurring in chaparral and expected to be drought-tolerant (Pavlik *et al*., 1991; Nixon, 2002). Within section *Lobatae*, the red oaks, we selected *Q. kelloggii*, a deciduous tree occurring in higher elevation woodlands that is expected to be drought-sensitive, and *Q. agrifolia*, an evergreen tree occurring widely along the California coast at lower elevations than *Q. kelloggii,* and expected to be more drought tolerant (Pavlik *et al*., 1991; Nixon, 2002).

Within section *Protobalanus*, we selected *Q. palmeri*, an evergreen shrub growing in dry regions like the Mojave Desert, as well as in Arizona and Baja California, expected to be a drought-tolerant species. *Q. chrysolepis*, an evergreen tree (sometimes growing as a shrub) that occurs in canyons, chaparral, and woodlands up to 2750 m in elevation is expected to be less drought-tolerant than *Q. palmeri* (Pavlik *et al*., 1991; Nixon, 2002).

### Sampling for functional traits

Individuals from each species were grown in a common garden at the California Botanic Garden (formerly Rancho Santa Ana) in Claremont, California (34.110738, −117.713913; 507 mm of rainfall per year; WorldClim, Fick & Hijmans, 2017). This common garden approach is ideal to test hypotheses related to trait evolution across species, since the observed differences represent heritable interspecies variation rather than phenotypic plasticity across multiple environments (Cordell *et al*., 1998; Dunbar-Co *et al*., 2009; Givnish & Montgomery, 2014; Mason & Donovan, 2015; Fletcher *et al*., 2018; Cavender-Bares *et al*., 2020). Data were collected from three to six individuals per species, based on availability in the common garden. For each individual, we used pole pruners to collect the most sun-exposed mature branch containing leaves grown in the current year, with no signs of damage and herbivory. Branches were carried to the lab in dark bags with moist paper and rehydrated overnight under plastic before harvesting stem sections and fully expanded leaves and stems for all subsequent analyses.

### Functional trait measurements

Functional traits were measured for three sun-exposed leaves per individual, unless noted otherwise. Leaf area was measured using a flatbed scanner and analyzed using the software ImageJ (imagej.nih.gov/ij/). After scanning, leaves were oven-dried at 70° for 72 h and dry mass was measured using an analytical balance (0.01 mg; XS205; Mettler-Toledo, OH, USA). Leaf mass per area (LMA) was calculated as lamina dry mass divided by leaf area (Pérez-Harguindeguy *et al*., 2013).

The concentrations of leaf nitrogen per mass (N_mass_) and the carbon isotope ratio (d^13^C) were determined from oven-dried leaves by the Center for Stable Isotope Biogeochemistry from the University of California, Berkeley, by continuous flow dual isotope analysis (CHNOS Elemental Analyzer interfaced to an IsoPrime100 mass spectrometer). The carbon isotope *_δ_*13 *_C_* −*δ*^13^ *C* discrimination (D^13^C; ‰) was calculated as: 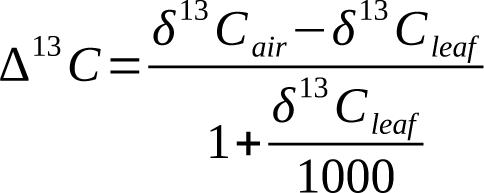, assuming a d^13^C_air_ of -8 ‰ *_δ_*13 *_C_* _1+_ *<u>leaf</u>* 1000 (Farquhar & Richards, 1984; NOAA Global Monitoring Laboratory 2018). The concentration of leaf phosphorus per mass (P_mass_) was determined from oven-dried leaves by the Baxter Lab in the Donald Danforth Plant Science Center, using a high throughput elemental composition (ionomics, approach from Salt *et al*., 2008, which uses Inductively Coupled Plasma-Mass Spectrometry (ICP-MS).

We estimated maximum rate of carboxylation per mass (V_cmax_) and electron transport rate (J_max_) from leaf N and P concentrations per mass (Domingues *et al*., 2010). Estimates of leaf lifetime integrated CO_2_ assimilation rate (A-_mass_) were derived from V_cmax_, J_max_ and isotope composition data using the Farquhar, von Caemmerer and Berry model (Franks *et al*., 2009).

We measured the wood density (WD) from 5-cm branch segments after bark removal using the water displacement method (Pérez-Harguindeguy *et al*., 2013). The osmotic pressure at turgor loss point (π_TLP_) was measured in two leaves rehydrated overnight from each of 5-6 studied individuals. We used vapor-pressure osmometers (Vapro 5520 and 5600, Wescor, US) to obtain the osmotic concentration at full turgor (π_o_) of the leaves and used calibration equations to estimate π_TLP_ (Bartlett *et al*., 2012). The leaf water potential at 50% loss of hydraulic conductivity (P50) was retrieved from Skelton *et al*. (2018), who performed the measurements using the optical method (Brodribb *et al*., 2016) in three branches from at least six individuals of seven of the 15 species included in this study grown in the Pepperwood Preserve in Sonoma County, located on the west coast of California (Skelton *et al*., 2018).

We measured leaf epidermal traits from microscopy images taken from nail varnish impressions of both leaf surfaces of one leaf from each sampled individual using the software ImageJ (http://imagej.nih.gov/ij/). We counted the number of stomata per image, measured the stomatal area (s), and calculated the stomatal density (d) as the number of stomata per area, the stomatal index (i) as the number of stomata per numbers of stomata plus epidermal pavement cells, and the maximum theoretical stomatal conductance (g_max_) as: *g max* =*bmds*^0.5^, where *b* ° *D*/*v*, *D* (2.55×10^-5^ m^2^ s^-1^) is the binary diffusivity of water vapor in air, *ν* (2.45×10^-2^ m^3^ mol^-1^) is the molar volume of air, and *m* is a dimensionless scaling factor representing conserved allometries among guard cell and stomatal pore dimensions (*m* ≍ 0.43 for kidney-bean shaped guard cells) (Franks et al 2009, Franks & Beeling 2009, Sack & Buckley 2016). Additionally, we used the minimum leaf conductance (conductance when stomata are fully closed, g_min_) as measured by Ochoa *et al*. (2024), measured using repeated gravimetric measurements (Scoffoni & Sack, 2010) for two leaves each from six individuals per species.

Principal component analyses (PCAs) were performed for two sets of traits. First, for traits which were measured on the same individuals used in the gene expression analysis, a PCA on all individuals was performed using LMA, WD, TLP, N_mass_, P_mass_, A_mass_, d, i, and g_max_. Second, a PCA was performed including additional drought-associated functional traits for which species means but not individual data were available: g_min_, P50_emb_, and D13C, in addition to TLP, LMA, and WD.

### Environmental variables for individual’s provenances and species’ native ranges

We obtained the coordinates for the location of sampling of acorns from the records of the California Botanic Garden and species occurrence data from the Global Biodiversity Information Facility (GBIF; references available in Table S2) and we used R software (version 3.4.4, R Core Team, 2019), to extract and calculate the mean, range and standard deviation of environmental variables of the range of distribution of each species. Occurrence records were downloaded using ‘rgbif’ and filtered to keep herbarium records only and remove incomplete (latitude or longitude missing) and duplicated records, non-natural occurrences (e.g., records from botanical gardens, planted urban trees) and to limit the temporal range to 1950-current (Riordan *et al*., 2015).

From open-access raster layers, we extracted environmental parameters: mean annual temperature (MAT), mean annual precipitation (MAP), mean temperature of the warmest quarter (T_warm_), temperature seasonality (T_s_), mean annual precipitation (MAP), precipitation of the driest quarter (P_dry_), precipitation of the warmest quarter (P_warm_; WorldClim, Fick & Hijmans, 2017), maximum temperature of the warmest month (T_max_), potential evapotranspiration (PET), aridity index (AI; CGIAR-CSI, NCAR-UCAR, (Zomer *et al*., 2008), soil water capacity (soil_WC_; ISRIC Soilgrids, (Hengl *et al*., 2017), length of the potential growing season (LPGS), precipitation of the potential growing season (GSppt), and mean temperature of the potential growing season (GStavg) (Table S2). The raster layers with same resolution were stacked using the stack function from the ‘raster’ package (Hijmans *et al*., 2020) (Hijmans, 2020) and the environmental parameters for each occurrence record were extracted using the extract function from the ‘dismo’ package (Hijmans *et al*., 2020). A PCA was performed on species mean climate variables, and climate PC axes were used to characterize multivariate species climate.

### Drying experiment

Branches were cut from the same adult individuals sampled for functional traits growing in a common environment at the California Botanic Garden, Claremont, California for use in the leaf drying experiment. Due to space constraints, the experiment was run on two dates (11 May and 22 May 2018), with individuals from the same species and accession spread across the two experiments to avoid batch effects. All conditions were kept the same between the two experiments. After collection, branches were re-cut under water and rehydrated overnight, and the treatment began the following day.

Two branchlets of approximately the same size (i.e., each having 8 leaves) were assigned to the control group or drought treatment. This allowed us to test the effects of drought on gene expression within the same individual and control for genetic variation in gene expression among individuals. Three replicates per treatment/species combination were used, including individuals from two accessions from a northern and southern part of the species range (except for *Q. durata*, which had one accession, and *Q. kelloggii*, which had three; Figure 1). Between 11 am and 12 pm, immediately after being cut, branches were re-cut underwater and placed on fans under high light to induce transpiration. Control branches were provided with water in a plastic bag wrapped around the base of the branch. This treatment continued for two hours, after which leaves were flash frozen in liquid nitrogen for RNAseq and stored at -80 °C (between 1 and 2 pm). Additionally, the leaf water potential (Ψ_leaf_) was measured for each branch before and after the experiment (pressure chamber; 0.001 MPa resolution, Plant Moisture Stress Model 1000; PMS Instruments Co), using one leaf before and two leaves after the experiment, to ensure controls were well-hydrated and to compare the degree of stress among the treated branches (Figure S7). For all species, Ψ_leaf_ values were near zero for the control leaves (species averages ranging from -0.04 to -0.09 MPa), and Ψ_leaf_ values decreased under the drying treatment as expected (species averages ranging from -0.76 to -3.26 MPa).

### RNA extraction and sequencing

To extract RNA from leaves, about 50 mg of leaf tissue was ground at room temperature using a Retsch mixer mill (model MM 301), placing grinding adapters and tubes in liquid nitrogen between rounds of grinding. For each extraction, tissue from three different leaves was used to reduce the effects of any variation among leaves within an individual branch. Polyphenolics and polysaccharides, which are common in oak leaves and interfere with downstream applications, were removed from leaves using a lithium chloride/urea-based pre-wash, as described in Mead *et al*. (2019). Tissue was kept in the buffer overnight at 4 °C. The next day, mRNA was extracted using the Sigma-Aldrich Spectrum Plant Total RNA Kit.

Sequencing libraries were prepared using the Illumina TruSeq RNA Library Prep Kit. RNA extractions were checked for quality using a NanoDrop instrument and Agilent TapeStation 2200, and final libraries were also quality checked on the Tapestation. Each library was diluted to 10 nM in EB buffer and 0.1% tween 20, based on DNA concentrations given by a Qubit Fluorometer, then samples were pooled by equal volume for sequencing. Samples were sequenced by the UCLA Broad Stem Cell Research Center’s sequencing core on an Illumina 4000 machine, using 100 bp, paired-end sequencing. Samples were sequenced across 3 lanes (12 libraries per lane) with samples from the same species and treatment date distributed evenly across different lanes; control and treated samples of the same individual were run on the same lane.

### RNAseq data processing

Sequence data files were converted from qseq to fastq format, demultiplexed (allowing for a one-base barcode mismatch), and reads failing the Illumina quality filter were removed using custom scripts (available at github.com/alaynamead/RNAseq_scripts). Adapters and low-quality reads (using a quality score cutoff of 27) were trimmed using Cutadapt 2.3. Reads were aligned to the *Quercus lobata* genome version 3.0 (Sork *et al*., 2022) using STAR 2.7 (Dobin *et al*., 2013) with default parameters. Only uniquely mapped reads (MAPQ score = 255) were retained. Sequencing platform duplicates (tagged ‘DT:Z:SQ’) were marked using the MarkDuplicates tool in Picard 2.18.23 and were removed. Read counts for each gene were obtained using the htseq-count command from HTSeq 0.11.2.

### Gene expression analysis

To identify differentially expressed (DE) genes, we used DESeq2 (Love *et al*., 2014) because it has high power for experiments with low sample sizes (Schurch *et al*., 2016). Only genes from chromosomes 1-12 were included in the analysis, excluding those from unplaced scaffolds, leaving a total of 38,510 genes used in the analysis. DE genes were identified using a species × treatment model, taking the paired design into account by including the individual as a factor. *P-*values were adjusted using the Benjamini-Hochberg procedure (Benjamini & Hochberg, 1995, FDR = 0.05).

We compared the gene expression responses across species to determine the factors shaping the evolution of the response to drought. To explore whether species were most similar to a closely related but ecologically different species, or to ecologically similar species from a different taxonomic section, we assigned species into ecologically-similar groups based on their drought tolerance assessed using π_TLP_, leaf habit, growth habit, and climatic niche. The deciduous trees, *Q. kelloggii* and *Q. lobata*, are the most drought-sensitive species. The other four species are all evergreen and relatively drought-tolerant, and we separated them into groups based on growth form: evergreen trees, *Q. agrifolia* and *Q. chrysolepis*; and evergreen shrubs, *Q. durata* and *Q. palmeri*. For each species, we compared its gene expression response to drying with the same-section species, and to the ecologically-similar species. The overlap in gene expression response to drying among species pairs was determined using the Jaccard index as in Bailey *et al. (*2015), calculated as the intersection (number of shared DE genes) divided by the union (total number of DE genes for both species). The same calculation was performed for the overlap among species of GO terms enriched within the DE genes.

To test whether patterns of gene expression were affected by alignment to the *Q. lobata* genome (because species more closely related to *Q. lobata* may align to the genome better than divergent species), we also aligned reads to the *Q. suber* genome (Ramos *et al*., 2018), which is within section *Cerris* and is an outgroup to all the species included in this study (Hipp *et al*., 2020). Alignment and differential expression analysis were done using the same methods as in the primary analysis.

For analyses of gene expression variation among individuals (described below), DESeq2 was used to produce normalized gene expression values from raw transcript counts. First, genes that were not expressed in this dataset were removed, leaving 36,331 genes. The ‘varianceStabilizingTransformation’ function with a parametric fit was used to transform count data, normalizing by library size across samples and stabilizing variance across genes (in particular, reducing the variance of lowly expressed genes) to better compare genes with different expression levels (Love *et al*., 2014).

GOseq (Young *et al*., 2010) was used to test whether the groups of upregulated and downregulated genes in each species were enriched for specific gene ontology (GO) terms. This method accounts for bias in differential expression results that are due to gene length, since longer genes will produce more reads during sequencing, resulting in greater statistical power for detecting expression differences. The default Wallenius distribution method was used to approximate the null distribution and calculate *P-*values. Each GO term is tested for both overrepresentation and underrepresentation, so *P*-values were adjusted separately for each group (using the Benjamini-Hochberg procedure) by determining whether a given GO term was more likely to be over- or underrepresented (based on *P*-values).

To test whether species differences were due to baseline gene expression in addition to gene expression plasticity, we identified genes for each pair that may be constitutively upregulated, or frontloaded (Barshis *et al*., 2013). Frontloaded genes in a given species (species 1) compared to another species (species 2) were defined as genes that were upregulated in response to drying in species 1, not differentially expressed in species 2, and having a higher expression under control conditions in species 2 than in species 1 (using a cutoff of 1.5, i.e. ≥1.5 times more highly expressed in species 2 under control conditions). To test the hypothesis that a given species had genes that were frontloaded compared to a second species, we tested whether frontloaded genes tended to be more highly expressed under control conditions in the second species, as might be expected if the gene is highly expressed at all times rather than plastically responding to a stimulus. For each species pair, the gene expression ratio under control conditions was calculated for the genes in the set of interest (genes that are upregulated in species 1 and non-DE in species 2), and the average ratio was compared to 10,000 replicates of a set of null genes (randomly sampled from all other genes). The *P*-value was calculated using the proportion of replicates in which the null set had a higher expression ratio than the test set.

We tested for relationships between gene expression response and drought tolerance traits in two ways. First, we tested for correlations between the magnitude of the species gene expression response (the number of DE genes) and potential explanatory factors: drought tolerance traits, species climate, and trait and climate PC axes. Second, we tested for correlations between drought tolerance traits and average expression of genes sharing the same GO term, which should be functionally similar. The average expression response for a given GO term was calculated as the individual difference in drying and control expression for each gene, averaged across all genes annotated with a given GO term. This difference was correlated with species average trait values to test for a relationship between expression response of similar genes and species drought tolerance traits. This analysis was done for a subset of GO terms likely to respond to drought: those that were significantly enriched in the DE genes, as well as terms related to abscisic acid, response to water/osmotic stress, oxidative stress, and photosynthesis, which are known to be physiological drought responses (Pinheiro & Chaves, 2011).

We used a partial redundancy analysis (RDA) in the R package vegan (Oksanen *et al*., 2019) to partition variance in gene expression response by the effects of phylogenetic relatedness and species climate. A redundancy analysis is a multivariate constrained ordination method that can be used to partition the variation in a data matrix into the proportion explained by two separate explanatory data matrices (Borcard *et al*., 2011). The gene expression response was measured as the difference in expression levels between drying and control branches within an individual. Pairwise branch distances were calculated for each species pair based on the oak phylogeny from Hipp *et al*. (2020) using the function ‘cophenetic.phylo’ from the R package ape, and the first and second axes of a PCoA on the branch distance matrix were used as variables measuring phylogenetic relatedness among species. The mean species climate was used as a measure of the climate a species has historically adapted to. We chose climate variables relevant to drought (precipitation of the driest quarter, mean temperature of the warmest quarter, potential evapotranspiration, soil water capacity, and temperature seasonality) and chose a subset of variables with low correlations with each other and that minimized variance inflation factors (tested using the function ‘vif.cca’). The final model included three variables: precipitation of the driest quarter, mean temperature of the warmest quarter, and soil water capacity. We used a series of models to determine how much each group of variables explained variation in gene expression: both climate and phylogeny, each factor alone, and each factor controlling for the effects of the other factor. We used permutation tests (running the ‘anova.cca’ function with 1000 permutations) to determine whether each model explained more variation in gene expression than expected by chance.

Models were run for two sets of genes: one set including all genes, and one set including the genes more highly ranked in the DE analysis. Because the number of significant genes varied widely among species, we did not choose these top genes by significance level to avoid species bias; instead, the top 500 genes from each species were selected as genes that are most likely to be plastic. Because of overlap in the top genes among species, this left 2078 “top” genes.

### Evolutionary divergence in gene expression

To identify genes with highly divergent gene expression among species, we performed a phylogenetic ANOVA using the expression variance and evolution (EVE) model (Rohlfs & Nielsen, 2015). This test uses the ratio of within-to between-species variance in gene expression (β), to identify genes that have high plasticity (high β) and genes that have high variation among species, but low variation within species (low β), consistent with directional selection on the expression levels of these genes. A likelihood ratio test (LRT) is used to determine whether a given gene likely has a different value of β than the rest of the genes. Only genes with no missing data were used (read count = 0 in one or more individuals), leaving 16,736 genes.

Significance was tested for each gene using 100 bootstrap replicates to calculate a null distribution of the likelihood ratio test statistic. This distribution simulated the gene expression using the maximum likelihood distributions under the null hypothesis (that β is the same across all genes). *P*-values were calculated by testing how many bootstrap replicates had a LRT value lower than the actual value. Low *P*-values indicate that the actual LRT is lower than expected under the null hypothesis, consistent with directional selection on gene expression that diverged among species. To identify genes that have diverged among drought tolerant and drought sensitive species, we took the genes with the top 1% LRT (169 genes, all had *P*-values of 0 or 0.01, meaning 0 or 1 bootstrap replicates had LRT values lower than the actual value). For these top genes, we tested for a correlation between the species average gene expression and π_TLP_.

## Results

### Variation across species in functional traits

Our experimental design paired species within sections with different climate niches and which did not necessarily share similar traits (Figure 1A). Thus, species did not cluster by section in a principal component analysis of species’ traits (Figure 1B). PC1 clearly separated the two deciduous and most drought-sensitive species, *Q. kelloggii* (section *Lobatae*) and *Q. lobata* (section *Quercus*) from the four evergreen species. Species from section *Protobalanus* (*Q. chrysolepis* and *Q. palmeri*), however, did cluster closely along PC1, perhaps because this clade is small and geographically restricted, consisting of only five evergreen species (Nixon, 2002). The designated drought-tolerant species within each section had a more negative π_TLP_, i.e., a lower wilting point. However, the difference in drought tolerance between species in the same section varied across the sections, with *Q. lobata* and *Q. durata* in section *Quercus* having strongly contrasting drought tolerance traits, whereas *Q. chrysolepis* and *Q. palmeri* in section *Protobalanus* were both relatively drought tolerant (Figure1B, Table S4).

### Water potential response to leaf drying experiment

Leaf water potential (Ψ_leaf_), an indicator of osmotic stress, was near 0 for all individuals in the control treatment, indicating leaves were well-hydrated (Figure S7). Under the drying treatment, all individuals had a negative Ψ_leaf_, indicating increased osmotic stress, ranging from -0.54 to -3.7 Mpa (Figure S7). Each species exhibited similar levels of stress except for *Q. palmeri*, which had a Ψ_leaf_ significantly greater (less negative) than all other species (Figure S7, pairwise t-test, p<0.05), likely because it is the most drought tolerant species in this study.

### Gene expression response to leaf drying experiment

Of the 38,510 annotated chromosomal genes, 36,331 were expressed in the leaf tissue. Species varied widely in the number of genes that were significantly differentially expressed between the control and drying treatments, with *Q. lobata* responding to drying with the most genes (5000, 13.8% of all genes) and *Q. palmeri* with the fewest (70, 0.2% of genes) (Table S1).

Across species, the oak species with traits adapted to aridity typically occurred in areas with higher potential evapotranspiration, and those species responded to leaf dehydration with fewer genes (Figure 2). Thus, across species, the number of differentially expressed genes increased exponentially in species with higher π_TLP_ values (Upregulated: *P* = 0.0001, R^2^ = 0.98; Downregulated: *P* = 0.0005, R^2^ = 0.95) and with the species-mean trait PC1 (Figure S1), which explains variation in the leaf water potential at 50% loss of hydraulic conductivity (P50_emb_), leaf potential at turgor loss point (π_TLP_), leaf mass per area (LMA), and wood density (WD) (Figure S5, *P* = 0.0043, R^2^ = 0.96). The number of DE genes was not associated with leaf water status (Ψ_leaf_) (Figure S6) (lm function in R, *P* >0.1). The results for the data aligned to the *Q. lobata* and *Q. suber* genomes were similar (Table S1), so further results reported here used the alignment to the chromosome-level *Q. lobata* genome rather than the scaffold-level genome of *Q. suber*.

**Figure 2.**
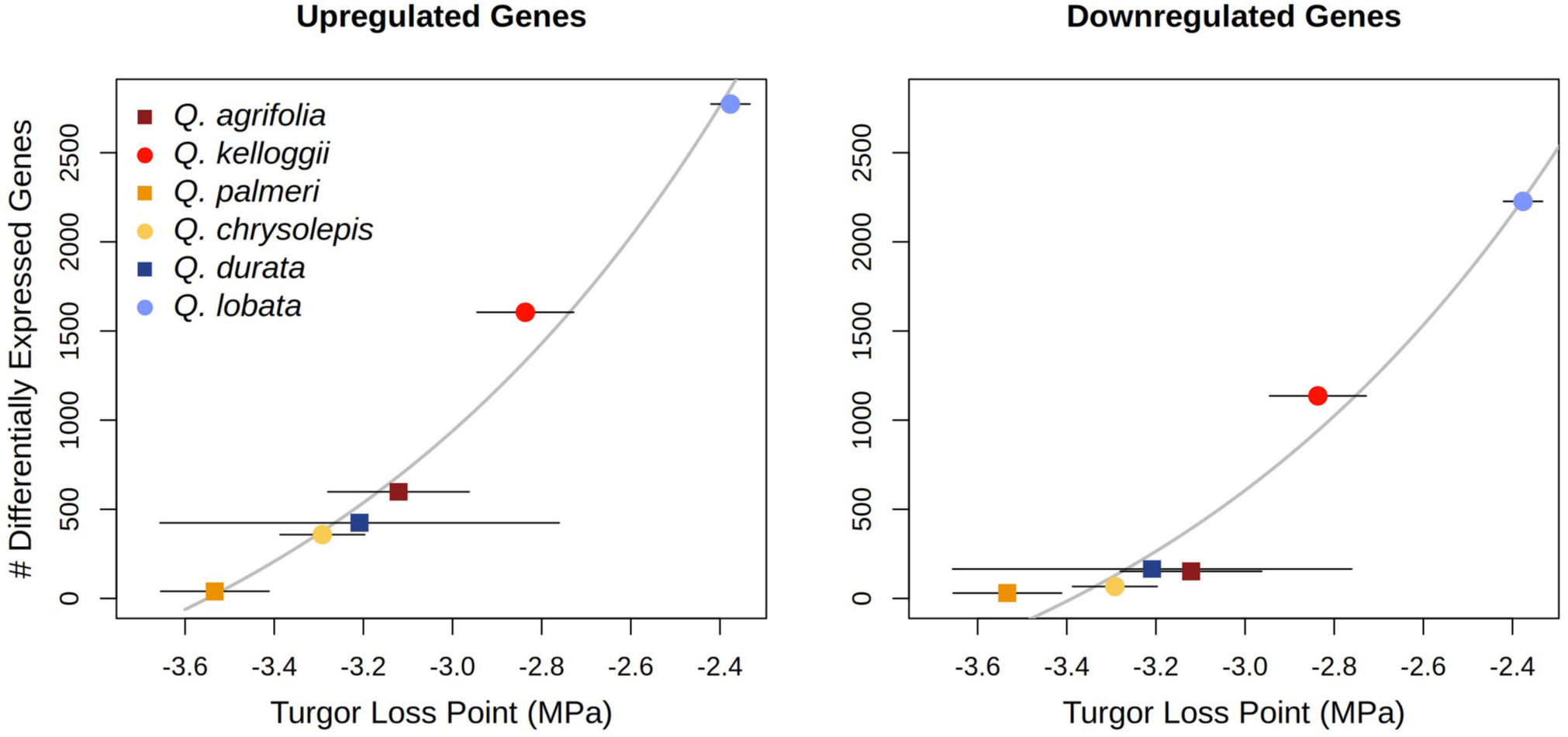
Relationship between plastic gene expression quantified as the number of differentially expressed genes and species average turgor loss point, the water potential at which a leaf loses turgor; higher (less negative) values indicate greater drought sensitivity, with upregulated and downregulated genes plotted separately. Error bars indicate standard error of turgor loss point; the number of differentially expressed genes was determined for each species rather than for individuals, so error bars are not shown. Grey line indicates an exponential fit (Upregulated: R^2^ = 0.98; *P* = 0.0001; Downregulated: R^2^ = 0.95; *P* = 0.0005).

Based on the GO enrichment analysis of differentially expressed genes, we identified GO terms that were significantly enriched within the upregulated and downregulated genes for each species (Table S6). Some GO terms were enriched within the upregulated genes for most species, except for one or both of the most drought-tolerant (*Q. palmeri* and *Q. chrysolepis*), including “calmodulin binding”, “DNA-binding transcription factor activity”, “protein ubiquitination”, and “response to water” (Table S6). Significantly enriched downregulated GO terms were more species-specific; however, multiple species downregulated “ribosome” and “translation” functions.

We identified several GO terms whose overall expression was correlated with drought tolerance traits across species. Of the traits tested, π_TLP_, an index of drought tolerance, had the greatest number of significant correlations with gene expression responses, followed by WD, leaf nitrogen content per mass (N_mass_), time-integrated photosynthetic rate per mass (A_mass_), and leaf phosphorus content per mass (P_mass_). For most of these GO terms, the species with the more drought-tolerant traits had little difference in expression between treatments, and the species with drought-sensitive traits more strongly upregulated or downregulated average GO expression in response to drying. More drought-sensitive species with higher π_TLP_ altered the expression of genes related to signaling (upregulated “double-stranded DNA binding” and “signal transduction”, downregulated “ubiquitin-protein transferase activity” and “sequence-specific DNA binding”). They also upregulated genes related to lipid metabolic process, which may be related to the reorganization of cell membranes that may occur during drought (Hoekstra *et al*., 2001; Gasulla *et al*., 2013); and upregulated genes involved in carbon metabolism, which may be related to the accumulation of solutes as a stress response, or may be the result of metabolism changes due to decreased photosynthesis rates (Pinheiro & Chaves, 2011) (e.g., processes involved in glycolysis, “beta-amylase activity”, and “trehalose biosynthetic process”). Drought-sensitive species with lower WD generally downregulated expression of ribosomal and translation genes, while higher WD species had more constant levels of expression of these genes, suggesting drought-tolerant species were able to maintain protein synthesis during the drying treatment, as found in drought-tolerant accessions of *Arabidopsis thaliana* (Des Marais *et al*., 2012) and populations of *Q. lobata* from warmer sites (Mead *et al*., 2019). Species with low WD also upregulated signaling-related genes, such as “MAP kinase activity” and “DNA binding”. Species with higher A_mass_ upregulated genes involved in “jasmonic acid biosynthetic process,” which can be a stress response (Wang *et al*., 2020).

### Comparisons of the gene expression response across species

The proportion of shared genes responding to drying for a species pair ranged from 22% (*Q. kelloggii* and *Q. lobata*) to just 1% (*Q. palmeri* compared to multiple species, Figure 2, Figure S2). We focused on testing whether species were more similar to the same-section species, as expected under neutral evolution, or to an ecologically similar species from a different section. At the level of individual genes, we found that four out of six species had gene expression responses that were more similar to the ecologically similar species than to the same-section species, consistent with parallel evolution of gene expression responses (Figure 3).

**Figure 3.**
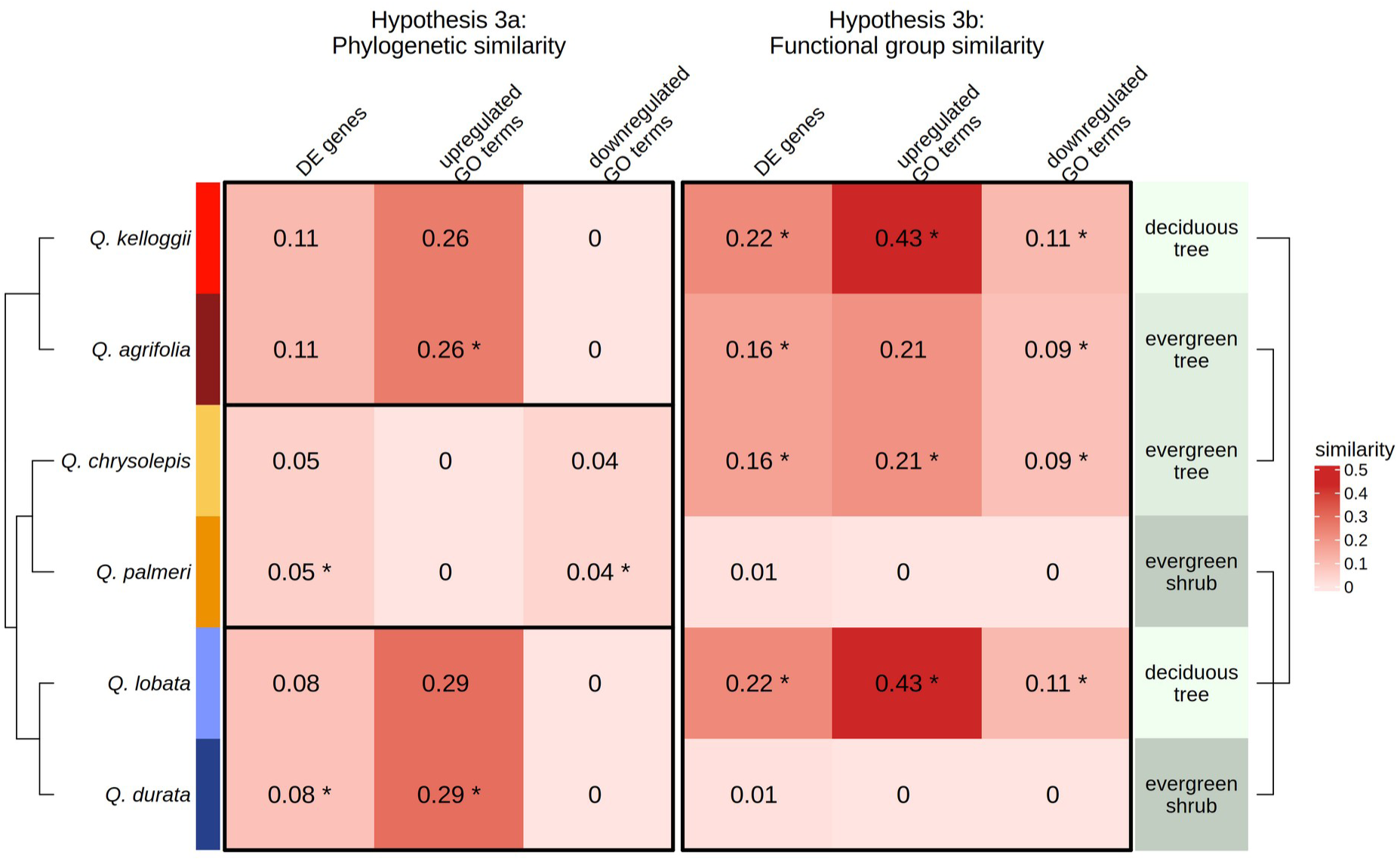
Heatmap indicating similarity between pairs of species for three factors under 2 different hypotheses: Hypothesis 3a, a species is more similar to its same-section species (i.e. within box for each species pair), as expected under neutral evolution, or Hypothesis 3b, more similar to its functionally similar species, as expected under parallel or convergent evolution in the response to drying stress (Hypothesis 2). Brackets on right indicate species pairs with same functional type that were compared to each other for Hypothesis 2. Similarity values for all possible species pairs are shown in Figure S2. Similarity was measured using the Jaccard index and calculates the overlap between two species (the intersection divided by the union) for either the genes that were significantly differentially expressed or for the GO terms that were significantly enriched in the set of upregulated or downregulated genes. A “*” indicates whether, for a given species and measure, it was more similar to the phylogenetically similar species (left); or the functionally similar species (right), suggesting parallelism or convergence.

We compared the differentially expressed genes to test which species pairs responded using genes with similar functional processes. Upregulated genes were generally more similar across species than downregulated genes; with the most similar species sharing 43% of their upregulated GO terms (Figure 3, Figure S2). Three species were more similar to the ecologically similar species than the same-section species, and of these species, similarity was higher for GO terms than for individual genes. However, similarity in the downregulated GO terms among species was lower; with the most similar species sharing only 11% of the downregulated GO terms. Four species had downregulated GO expression that was most similar to the ecologically similar species.

We expected species occurring in similar climates to have similar gene expression responses. However, we found that the similarity in gene expression response among two species was not related to the similarity in their average climate, and in fact some of the most similar species’ pairs came from contrasting climates (Figure S4). Species with similar responses did have relatively high overlap in their ranges (Figure S3); but the species with the highest overlap in ranges (*Q. agrifolia* and *Q. kelloggii*) were not particularly similar in their gene expression response (Figure S2). We used a redundancy analysis to further explore the relationship among functional traits, gene expression, climate, and phylogeny.

### Redundancy Analysis

The redundancy analysis partitioned the interspecific variation in functional traits and the gene expression response to drying into that explained by phylogeny alone, climate alone, or their joint influence (Table 1). We found that a species’ climate explained equally as much or more variation as did phylogeny for all the phenotype sets tested: functional traits, the response to drying across all genes, and the response to drying for the top DE genes. The overall amount of variation explained (R^2^ of the full model, Table 1) varied between the two gene expression sets, with climate and phylogeny explaining more variation in the response of the top DE genes than they did in the dataset containing all genes. The degree to which the influence of climate and phylogeny could be separated also varied among the functional traits and gene expression: the joint influence of climate and phylogeny explained little to none of the variation among species in gene expression, but explained 26% of the variation in functional traits.

**Table 1.** Redundancy analysis partitioning the effects of climate (species mean) and phylogeny (phylogenetic distance) on the variance in functional traits and gene expression difference in the control and branch-drying treatment for an individual plant. The joint influence of phylogeny and climate explains the portion of the variation that cannot be disentangled and is calculated by subtracting the variance explained by phylogeny alone and climate alone from the total variance explained by the full model; *P*-values are not given because it is not testable.

|  | Functional traits |  |  |  | All genes |  |  |  | Top DE genes |  |  |  |
| --- | --- | --- | --- | --- | --- | --- | --- | --- | --- | --- | --- | --- |
| | df | Adjusted $R^2$ | $p$ | | df | Adjusted $R^2$ | $p$ | | df | Adjusted $R^2$ | $p$ | |
| full model | 5 | 0.50 | 0.001 | *** | 5 | 0.09 | 0.001 | *** | 5 | 0.40 | 0.001 | *** |
| phylogeny alone | 2 | 0.11 | 0.016 | * | 2 | 0.04 | 0.035 | * | 2 | 0.15 | 0.001 | *** |
| climate alone | 3 | 0.12 | 0.030 | * | 3 | 0.06 | 0.008 | ** | 3 | 0.24 | 0.001 | *** |
| Joint influence of phylogeny and climate | - | 0.26 | - | - | - | 0 | - | - | - | 0.01 | - | - |

### Identification of frontloaded and evolutionarily diverged genes

We tested whether frontloaded genes, which are constitutively upregulated, could contribute to drought tolerance across oak species. We identified frontloaded genes as those that were upregulated in a given drought-sensitive species, non-DE in given drought-tolerant species, and more highly expressed in the drought-tolerant species under control conditions (Table S7). This produced a set of genes for each species pair that were constitutively highly expressed in one species in comparison to the other species. For example, 149 genes were upregulated as a drought response in *Q. lobata* (drought-sensitive) and were more highly expressed in *Q. palmeri* (drought-tolerant) under the non-stressful control conditions. Generally, there was little overlap among the gene sets for each species pair; however, we identified 8 genes that were frontloaded in the most drought-tolerant species *Q. palmeri* when compared to the two least drought-tolerant species, *Q. kelloggii* and *Q. lobata*. These genes were annotated with GO terms that included protein kinase activity, DNA-binding transcription factor activity, and metal ion binding.

Using the expression variance and evolution (EVE) model (Rohlfs & Nielsen, 2015), we identified genes with strong divergence in expression levels across species and then identified genes within this set that had expression levels correlated with drought tolerance across species (Figure 4, Table S8). Of the 169 genes in the top 1%, 26 had expression levels that were significantly correlated with drought tolerance (as measured with π_TLP_). Of these, 12 were more highly expressed in the drought-tolerant species, and 14 were more highly expressed in drought-sensitive species. Many of these correlations seemed to be driven by divergent expression in *Q. lobata*, the least drought-tolerant species; in Figure 4, we present correlations for which variation among all six species is more evident. This set of genes included several genes related to signaling had higher expression in the drought-tolerant species, including QL03p063179 (a gene annotated with the GO terms protein kinase activity; ATP binding; and protein phosphorylation), QL02p061158 (DNA-binding transcription factor activity and regulation of transcription, DNA-templated), and QL05p003384 (SAGA complex and transcription coregulator activity). These genes may enable drought tolerance through their higher baseline expression rather than a plastic response to drought. QL05p055903 (microtubule motor activity; microtubule-based movement; ATP binding; and microtubule binding) was one of the genes with higher expression in drought-sensitive species.

**Figure 4.**
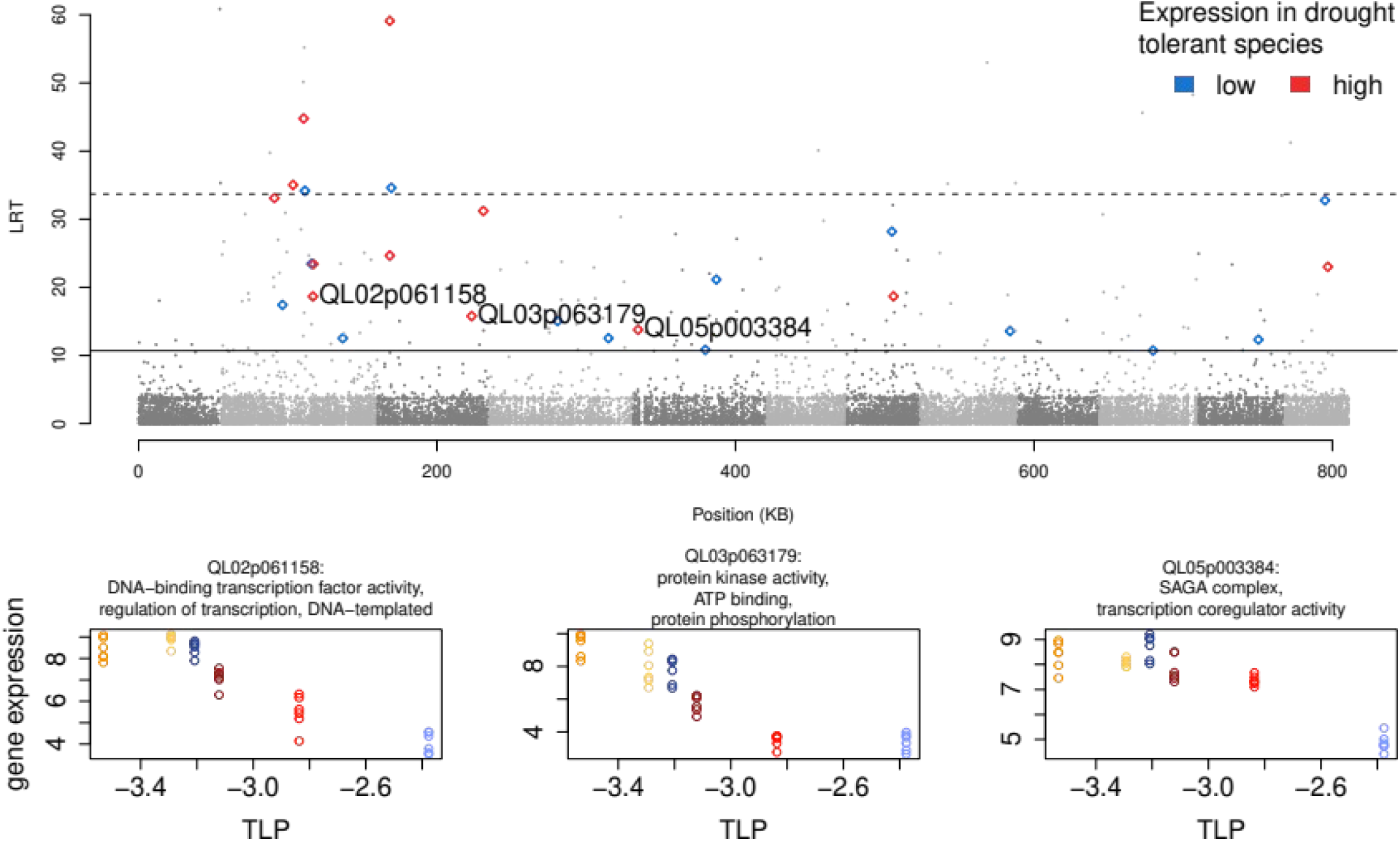
Manhattan plot with the likelihood ratio on the y-axis and genomic position on the x-axis. Each point is a gene; dark and light grey colors represent different chromosomes. Genes with higher likelihood ratio test (LRT) values have expression levels that are more divergent among species. Solid line represents the 99th percentile, dashed line represents the 99.9th percentile. Diamonds indicate genes that have LRT values in the top 1% and whose gene expression is correlated with drought tolerance (via turgor loss point). Blue diamonds have lower expression in more drought tolerant species and red diamonds have higher expression in more drought tolerant species. (B-D) Selected genes that are more highly expressed in drought-tolerant species. While some genes correlated with drought tolerance were more strongly differentiated among species (indicated by a higher LRT), many of these relationships were primarily driven by high or low expression in *Q. lobata* compared to all other species, so here we present genes for which variation among all six species is more evident. Titles give the gene name and associated GO terms.

## Discussion

This study finds clear patterns of constitutive and differential gene expression associated with drought tolerance traits across oak species. In response to a leaf drying treatment that simulates drought conditions, our findings demonstrate that the more drought-tolerant species exhibit high constitutive gene expression of drought-associated genes, while drought-sensitive species have greater gene expression plasticity, and that some of these genes and gene functions have repeatedly evolved similar expression patterns across oak sections separated by 45 million years of evolution.

### Drought-tolerant species have less plasticity but high levels of constitutive gene expression

Our comparison of gene expression in response to drought across California oak species reveals the novel finding that drought tolerance is associated with lower plasticity and higher constitutive gene expression, consistent with hypotheses 1 and 2. This pattern may arise because the functional traits of more drought-tolerant species allow them to withstand dehydration without significantly altering their gene expression, while drought-sensitive species must respond to drought through plasticity mediated by gene expression. Drought tolerance in these oak species may have evolved primarily through mechanisms besides phenotypic plasticity, such as mutations affecting protein-coding regions, or constitutive changes in gene expression, consistent with hypothesis 2. We identified two groups of genes whose gene expression patterns may contribute to drought tolerance through non-plastic mechanisms: genes that were frontloaded in drought-tolerant species and genes whose expression levels have evolutionarily diverged across species and were also associated with drought tolerance. These genes could underlie functional drought tolerance traits or reduce the effects of drought stress on leaf tissue, making an immediate plastic response unnecessary.

Within these two sets of genes, several had functions related to signaling. Signaling genes could be beneficial as frontloaded and/or constitutively highly expressed genes in drought-tolerant species, as they may control networks of genes with related functions, resulting in a large overall phenotypic effect (Des Marais & Juenger, 2010; Todaka *et al*., 2015, but see Des Marais *et al*., 2017). It is possible that the lack of plasticity in drought-tolerant species evolved from an ancestrally plastic state through canalization (Rivera *et al*., 2021), however, in this study we cannot infer the ancestral state of gene expression to evaluate when shifts may have occurred.

The limited number of previous studies testing whether differential gene expression under stress corresponds to stress tolerance across taxa do not reveal a general trend (Sandoval-Castillo *et al*., 2020; Bittner *et al*., 2021; Komoroske *et al*., 2021; Roelofs *et al*., 2009; Steele *et al*., 2025). Similar to our findings, the drought tolerant oak species *Q. douglasii* exhibited constitutive upregulation rather than a plastic response to drought stress (Steele *et al*., 2025).

However, the trends differ in animal experiments. One study in house mice found that a desert population responded to dehydration with fewer differentially expressed genes compared to a non-desert population, and the gene expression of non-desert mice became more similar to desert mice under water stress, indicative of adaptive plasticity in response to dehydration (Bittner *et al*., 2021). However, several studies in fish have found that heat-tolerant species had higher gene expression plasticity in response to heat stress than heat-sensitive species (Sandoval-Castillo *et al*., 2020; Komoroske *et al*., 2021). While the degree of expression plasticity measured in different taxa was likely influenced by the specific experimental conditions, it may also indicate variation in the stress tolerance mechanisms evolved by the taxa. For increased plasticity to be beneficial, the organism must be able to accurately detect and respond to the environmental stress with minimal lag time, and there must be a cost to expressing the stress-induced phenotype under benign conditions (DeWitt *et al*., 1998). If these conditions are not met, stress tolerance is more likely to evolve through other adaptations. Less gene expression plasticity in the four most drought-tolerant and evergreen oak species may reflect investment in drought-tolerant leaves, which have a high carbon cost and must survive multiple years of summer drought. The two more drought-sensitive deciduous species regenerate their leaves each year, so the carbon cost of more drought-tolerant leaves may outweigh the cost of gene expression plasticity in wetter environments.

### Climate niche influences species differences in functional traits and responses

Across the six oak species, climate explained more of the variation in plastic gene expression in response to leaf drying than phylogeny (Table 1). Under neutral evolution or adaptation constrained by phylogenetic history, phylogeny would be expected to explain more interspecific variation than climate. Instead, our results suggest that closely related oak species diverged in their adaptation to contrasting climates by evolving traits and gene expression patterns beneficial to their climates. Further, both climate and phylogeny explained more variation in gene expression in the dataset containing the genes with the most plastic response to drying than they did in the dataset containing all genes, suggesting the drying-responsive genes are shaped by both neutral and adaptive evolution more strongly than non-DE genes.

Notably, while climate explained some of the variation in the multivariate gene expression response to drought across species, it did not explain pairwise species similarities. Thus, the species pairs with the most similar responses do not occur in similar climates (Figure S4). This finding contrasts with that of another study which found that tree species with similar gene expression responses to drought tended to co-occur within a forest plot (Swenson *et al*., 2017). Similarly, gene expression and trait PCs were not strongly correlated with climate PCs; the number of differentially expressed genes were most associated climate with PC4 (associated with higher PET and longer potential growing season), but not with precipitation or aridity index (Figure S5). The fact that two oak species occur in similar climates today does not mean that they will necessarily converge on the same traits; the relationship among the environment, traits, and fitness is complex and our results must be interpreted with an understanding of species’ biology and evolutionary history. While species with shared responses to leaf drying did not typically occur in similar climates, they did often have similar functional traits. Deciduousness can mediate drought tolerance strategies (Kaproth *et al*., 2023), and is one factor that may explain variation observed in our study species. Species’ functional traits may be more important in determining their transcriptomic drought response than climate because traits can modulate the stress experienced at the tissue and cell level for a given climate, and there are multiple possible combinations of traits allowing adaptation to the same conditions (Sack & Buckley, 2020). For example, *Q. agrifolia* and *Q. lobata* occur in similar climates, often at the same sites, but have contrasting traits and gene expression responses. *Q. lobata* has more drought-sensitive traits (Figure 1B) despite occurring in relatively hot, dry climates, possibly because it is adapted to cooler conditions than it experiences in its current range (Browne *et al*., 2019), and it survives in these dry regions by colonizing microhabitats such as valleys where groundwater reserves can be accessed with its deep roots (Mahall *et al*., 2009). These characteristics of the deciduous *Q. lobata* may indicate a drought avoidance strategy, while the evergreen *Q. agrifolia* may be better able to tolerate lower water potentials. Indeed, it is common for co-occurring oaks in western North America to have contrasting leaf traits, potentially allowing co-existence through complementary resource acquisition strategies (Cavender-Bares *et al*., 2018).

### Parallelism and convergence in gene expression responses to leaf drying

We found evidence of parallelism, defined as use of the same genes, in the transcriptomic response to drought in four of six ecologically-similar but evolutionarily diverged species: all four tree species shared more differentially expressed (DE) genes with their distantly-related, ecologically similar species than with a closely related species (Figure 3). These results suggest that adaptive gene expression responses contribute to divergence in drought tolerance across the oak phylogeny, supporting hypothesis 3. This finding could result from more rapid divergence of gene expression patterns than protein-coding mutations because they have fewer negative pleiotropic effects. Thus, our findings imply that plastic gene expression was decoupled from phylogenetic structure and, instead, it is more tightly linked to a species’ climate niche.

We tested whether species showed gene expression patterns more similar to a close relative or to an ecologically-similar species. Notably, the species with the most similar plastic gene expression response were the least drought-tolerant species pair, the deciduous trees; followed by the moderately drought-tolerant evergreen trees (Figure 3, Figure S2A). This finding supports hypothesis 3; suggesting that natural selection has acted similarly on gene expression responses because these four tree species independently adapted to similar conditions. However, the drought-tolerant shrub species *(Q. durata* and *Q. palmeri)* did not show evidence of a parallel response and were more similar to a closely-related species than to each other. If gene expression plasticity is not a primary contributor to drought tolerance in these species, a lack of parallel responses may simply result from reduced selection pressure on the gene expression response to drought. Overall, parallel selection on gene expression was supported in oaks, but the majority of genes used in drought response are species-specific.

Additionally, the level of parallelism in gene expression identified in this study (16-22%) is lower than that found in two conifer species, where 74% of the orthologs with plastic responses showed similar expression patterns in both species (Yeaman *et al*., 2014). However, a study on two *Eucalyptus* species found limited population-level parallelism in gene expression (Ahrens *et al*., 2022). Higher parallelism in conifers may result from slower divergence in gene expression divergence compared to angiosperms (Walia *et al*., 2009; Yeaman *et al*., 2014; Bouzid *et al*., 2019). Overall, our results suggest both parallel (same-gene) and convergent (same-function) evolution of gene expression responses in the four tree species, but not the drought-tolerant shrubs.

We also found evidence for convergence, or expression of different genes with similar functions, for the ecologically-similar species *Q. kelloggii* and *Q. lobata* (the deciduous trees) and *Q. chrysolepis* with *Q. agrifolia.* Two of the three ecologically-similar species pairs had greater overlap in their upregulated GO terms than individual genes (Figure 3), supporting the hypothesis that repeated evolution of traits is more likely to occur through evolution at higher organizational levels (Agrawal, 2017). Interestingly, the pair of deciduous tree species, which had the highest level of parallelism in DE genes, also showed the highest level of convergence compared to other species pairs. However, we also note that some species that were neither closely related nor functionally similar also showed high overlap in the types of genes that responded to drying, such as the similarity in upregulated GO terms between *Q. durata*, a drought-tolerant evergreen shrub, and *Q. kelloggii*, a drought-sensitive deciduous tree (Figure S2B), indicating that there are additional factors contributing to gene expression similarity. In contrast to the upregulated genes, genes that were downregulated in response to drying had fewer GO terms shared across species. In response to stress, plants typically upregulate drought-specific responses while downregulating their normal processes such as growth and photosynthesis (Chaves *et al*., 2003), so the lack of convergence in downregulated functions may be driven by species differences under non-stressful conditions. While we found evidence for convergence in upregulated GO terms associated with drought response and signaling, the enriched GO terms within the downregulated genes were usually species-specific and included terms related to photosynthesis and chloroplasts, transport, cell walls, and mitochondria (Tables S5, S6). These results suggest that oaks may evolve convergent responses to stress through evolutionary changes in different genes, consistent with population-level studies on oaks, which have found that populations with the same ecophysiological responses to drought used different gene networks in their drought response (Mead *et al*., 2019).

## Conclusions

Stress tolerance is a crucial aspect of evolutionary response to climate change. Unexpectedly, we discovered that drought tolerance in some oaks is facilitated by the evolution of constitutive patterns of gene expression rather than plasticity in gene expression triggered by the environment. We found that the most drought-tolerant species responded to leaf drying with fewer genes than the drought-sensitive species, demonstrating that short-term plasticity in leaf gene expression is not the primary mechanism of drought tolerance among these California oak species. Conversely, drought-sensitive oak species have a strong plastic response to drought, which has evolved through a combination of parallel changes in the same genes, convergent changes in genes with similar functions, and species-specific genes. Our results point to constitutive gene expression as a potentially important evolutionary target for selection on drought response in oaks.

## Data Availability

RNAseq data is available at the NCBI SRA (Accession: PRJNA1300321, https://www.ncbi.nlm.nih.gov/bioproject/1300321).

Trait and climate data are available as Supplementary Tables 2-4.

Analysis scripts for gene expression data are available at https://github.com/alaynamead/oak_comparative_drought_genomics

## Acknowledgements

We acknowledge the native peoples of California as the traditional caretakers of the oak ecosystems sampled for this project.

We thank the California Botanic Garden for allowing us to sample, and Alec Baird, Marvin Browne, Leila Fletcher, Christian Henry, Yao Li, Scott O’Donnell, and Santiago Trueba for helping with sampling and providing useful feedback on the manuscript.

Genomic analyses, sequencing costs, and salary for A.M. were supported by National Science Foundation Plant Genome Research Program awarded to VL Sork (NSF IOS-#1444661). C.M. was supported by the Brazilian National Research Council (CNPq) through the Brazilian Science Without Borders Program (grant number: 202813/2014-2).

