## Supplementary Information for "Drought tolerance is associated with constitutive gene expression, not plasticity, across California oak species"

#### Supplementary Results and Discussion

*Gene functions associated with drought tolerance*

There were multiple GO terms with expression levels correlated with species' drought traits. Because many of these traits were correlated with each other, for each GO term we identified the trait with the highest correlation. Most GO terms with significant trait correlations were best correlated with turgor loss point ($\pi_{TLP}$), followed by wood density (WD), leaf nitrogen content per mass (Nmass), time-integrated photosynthetic rate per mass (Amass), and leaf phosphorus content per mass (Pmass). For most of these GO terms, the species with the more drought-tolerant traits had little difference in expression between treatments, and the species with drought-sensitive traits more strongly upregulated or downregulated average GO expression in response to drying. More drought-sensitive species with higher $\pi_{TLP}$ altered the expression of genes related to signaling (upregulated "double-stranded DNA binding" and "signal transduction", downregulated "ubiquitin-protein transferase activity" and "sequence-specific DNA binding"). They also upregulated genes related to lipid metabolic process, which may be related to the reorganization of cell membranes that may occur during drought (1, 2); and upregulated genes involved in carbon metabolism, which may be related to the accumulation of solutes as a stress response, or may be the result of metabolism changes due to decreased photosynthesis rates (3) (e.g. processes involved in glycolysis, "beta-amylase activity", and "trehalose biosynthetic process"). Drought-sensitive species with lower WD generally downregulated expression of ribosomal and translation genes, while higher WD species had more constant levels of expression of these genes, suggesting drought-tolerant species maintained protein synthesis during the drying treatment, as found in drought-tolerant populations of *Q. lobata (4)* and drought-tolerant accessions of *Arabidopsis thaliana* (5). Low WD species also upregulated signaling-related genes, such as "MAP kinase activity" and "DNA binding". Species with higher Amass (photosynthetic rate) upregulated genes involved in "jasmonic acid biosynthetic process," which can be a stress response (6).

*Gene expression divergence*

We identified genes that diverged among drought-tolerant and drought-sensitive species regardless of their phylogenetic relatedness by testing for correlations between gene expression and $\pi_{TLP}$ among the genes with the strongest expression divergence among species. For example, several genes related to signaling had higher expression in the drought-tolerant species, including QL03p063179 (a gene annotated with the GO terms protein kinase activity; ATP binding; and protein phosphorylation) QL02p061158 (DNA-binding transcription factor activity and regulation of transcription, DNA-templated), and QL05p003384 (SAGA complex and transcription coregulator activity). As these genes have higher baseline expression rather than higher plasticity in drought-tolerant species, they may be drought adaptations that allow these species to have lower gene expression plasticity under drying stress. QL05p055903 (microtubule motor activity; microtubule-based movement; ATP binding; and microtubule binding) was one of the genes with higher expression in drought-sensitive species. Although the specific functions of these genes are unclear, they could theoretically relate to the trade-off between drought tolerance and faster growth under ideal conditions, and higher expression could be selected for or selected against under different levels of water availability (7).

**Table S1.** Number of differentially expressed (upregulated and downregulated) genes by species, for datasets aligned to the *Q. lobata* genome (primary analysis) and the *Q. suber* genome.

| | *Q. lobata* alignment upregulated | *Q. lobata* alignment downregulated | *Q. suber* alignment upregulated | *Q. suber* alignment downregulated |
|---|---|---|---|---|
| *Q. agrifolia* | 598 | 152 | 591 | 162 |
| *Q. chrysolepis* | 358 | 67 | 364 | 69 |
| *Q. durata* | 424 | 165 | 411 | 173 |
| *Q. kelloggii* | 1605 | 1136 | 1671 | 1134 |
| *Q. lobata* | 2773 | 2227 | 2796 | 2069 |
| *Q. palmeri* | 40 | 30 | 46 | 28 |

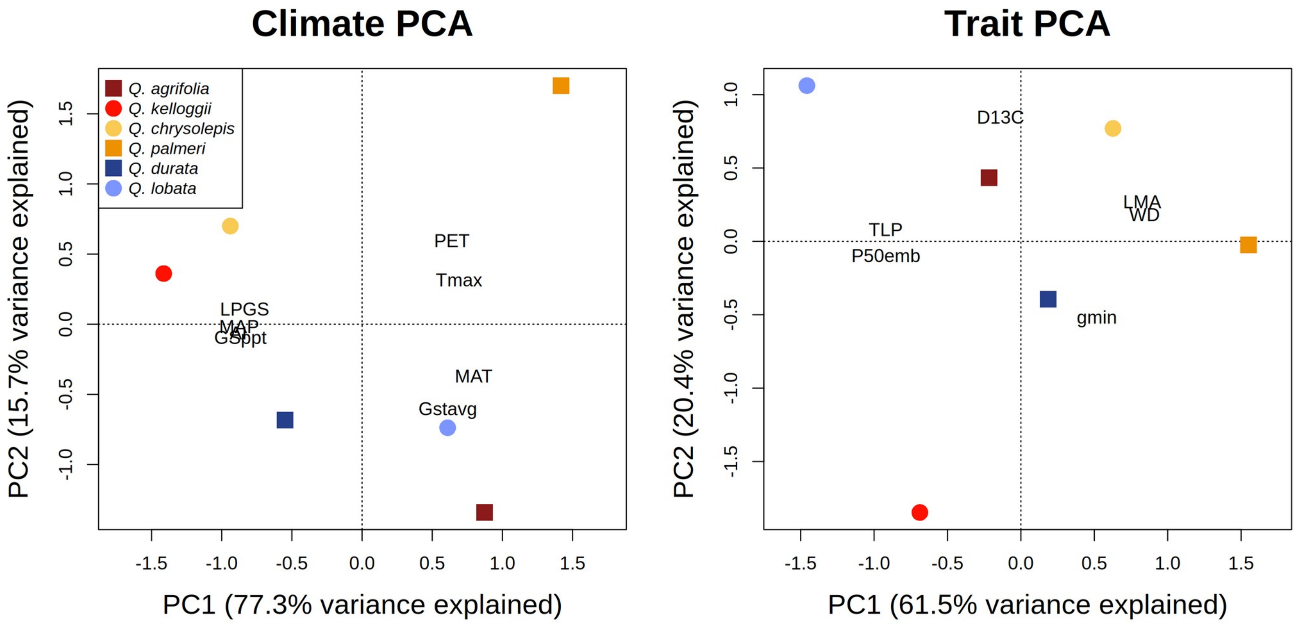


**Figure S1**. PCAs of species-mean climate variables and trait measurements, performed using the rda function from the vegan package (version 2.6-10), in R version 4.5.1. Climate variables include: mean annual temperature (MAT), mean annual precipitation (MAP), aridity index (AI; CGIAR-CSI, NCAR-UCAR, Zomer *et al.*, 2008), potential evapotranspiration (PET), length of the potential growing season (LPGS), precipitation of the potential growing season (Gsppt), and mean temperature of the potential growing season (Gstavg). Trait variables include leaf mass per area (LMA), wood density (WD), turgor loss point (TLP), carbon isotope discrimination (D13C), minimum stomatal conductance ($g_{min}$), and vulnerability to embolism ($P50_{emb}$).

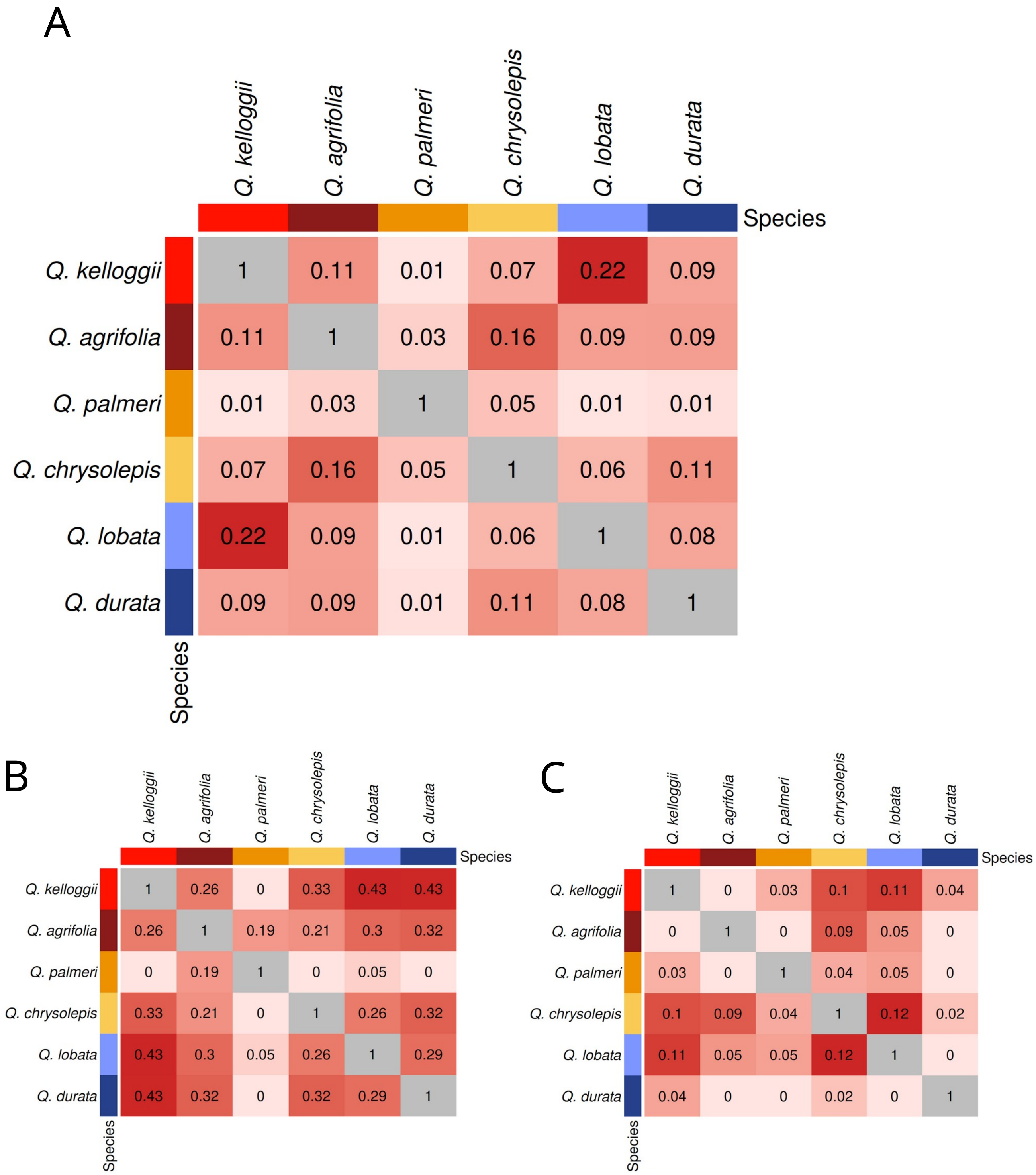


**Figure S2.** Overlap between all species pairs in (A) the significantly DE genes, and GO terms that were significantly enriched within the genes that were upregulated under drying (B) or downregulated (C).

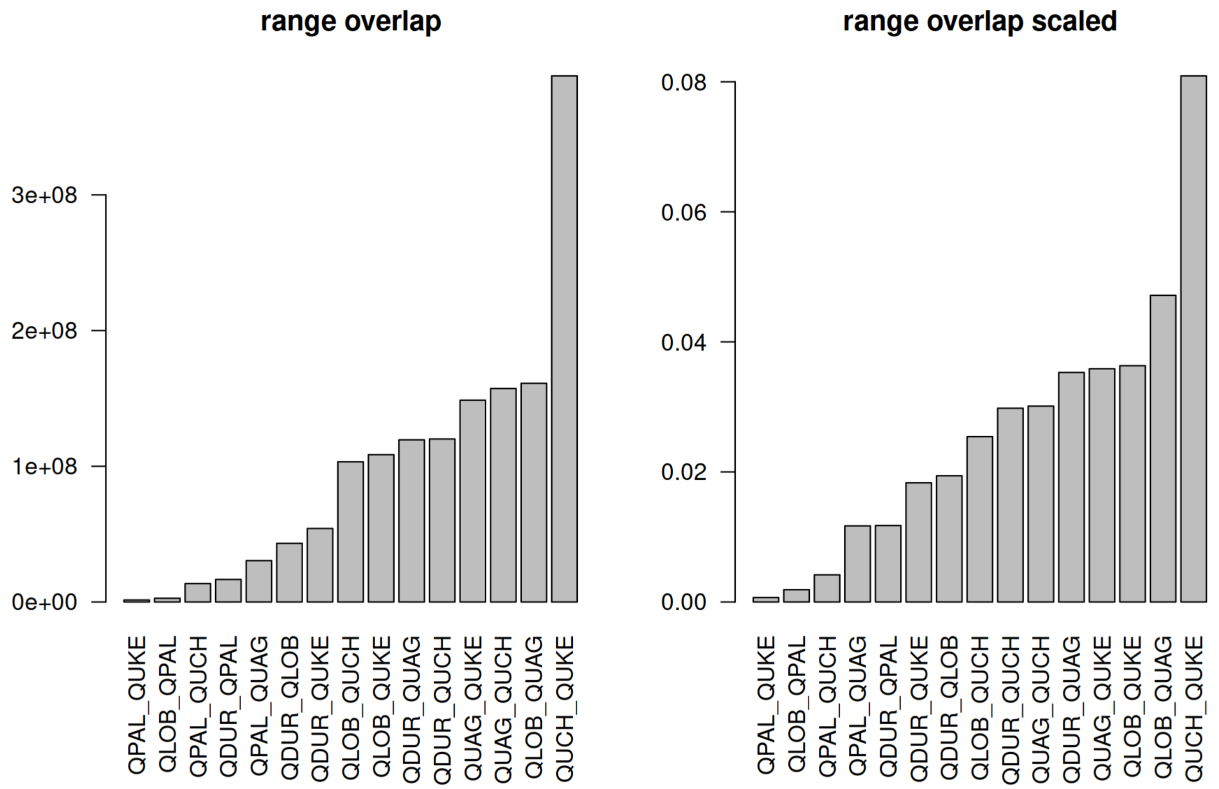


**Figure S3.** Overlap in species ranges calculated using a 1 km radius around occurrence points for each species. Right graph is the overall area overlap, left graph is scaled by the total area occupied by both species in the pair.

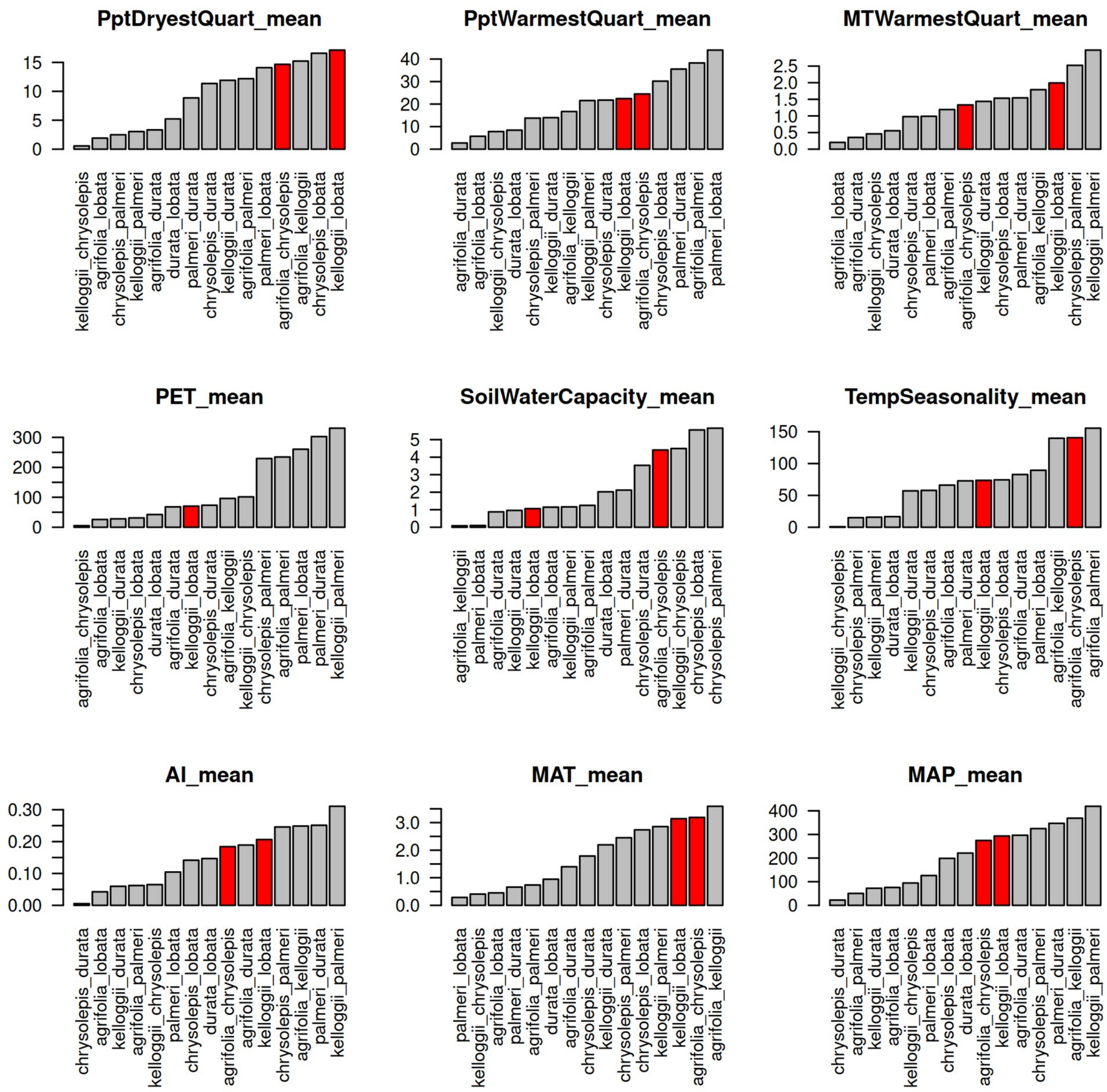


**Figure S4.** Climate difference among species pairs, calculated as the difference in mean climate values for each species. The two functionally similar species pairs which had greater similarity in gene-level expression plasticity than a same-section species, suggesting parallel evolution, are highlighted in red. Full names for climate variables are as follows: precipitation of the driest quarter, precipitation of the warmest quarter, mean temperature of the warmest quarter, potential evapotranspiration, soil water capacity, temperature seasonality, aridity index, mean annual temperature, and mean annual precipitation.

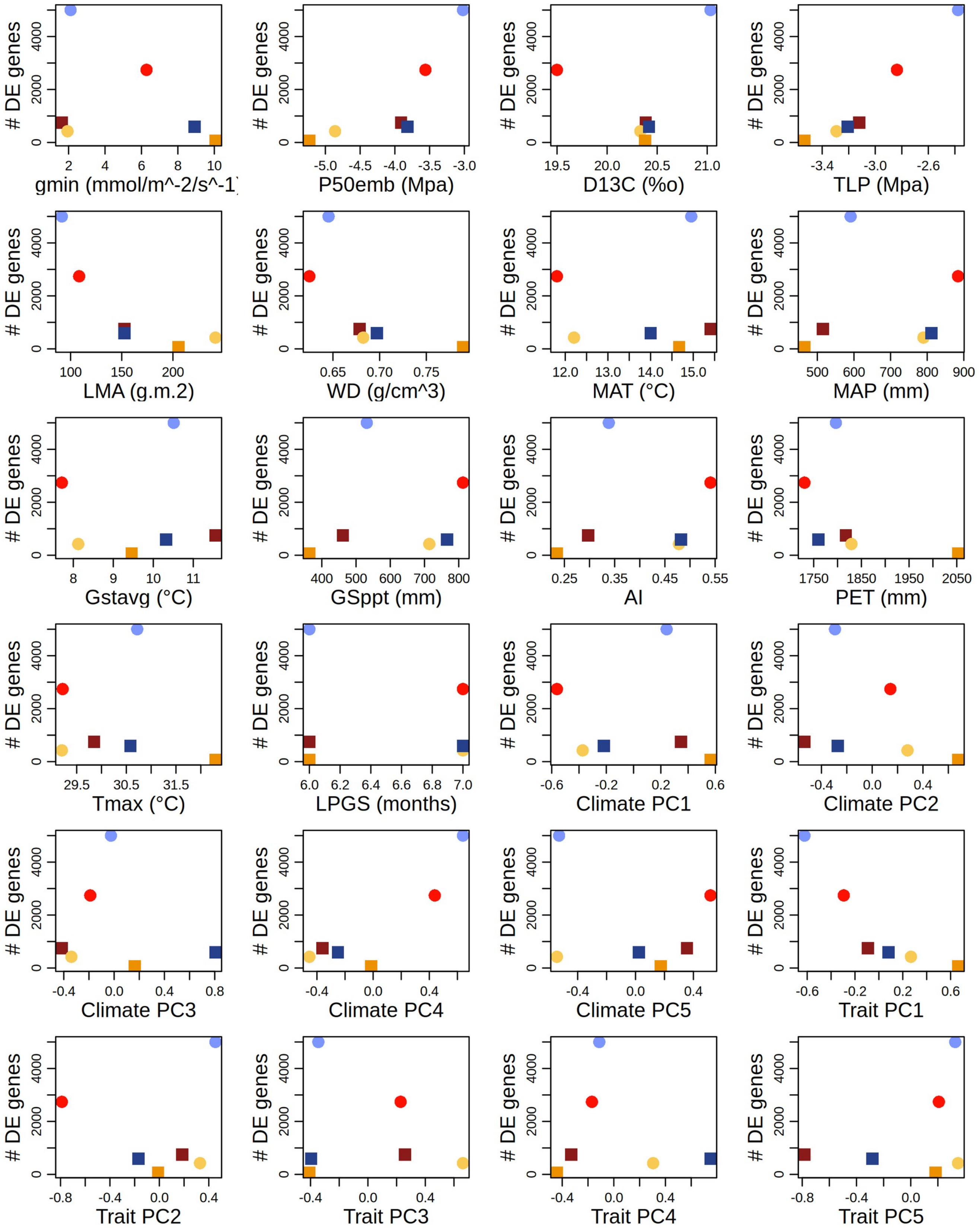
### DE genes
gmin (mmol/m^-2/s^-1)
P50emb (Mpa)
D13C (‰)
TLP (Mpa)
LMA (g.m.2)
WD (g/cm^3)
MAT (°C)
MAP (mm)
Gstavg (°C)
GSppt (mm)
AI
PET (mm)
Tmax (°C)
LPGS (months)
Climate PC1
Climate PC2
Climate PC3
Climate PC4
Climate PC5
Trait PC1
Trait PC2
Trait PC3
Trait PC4
Trait PC5

**Figure S5.** The relationship between the number of differentially expressed genes  and species average values of drought-associated functional traits and climate variables. Trait abbreviations are: leaf mass per area (LMA), wood density (WD), turgor loss point (TLP), leaf nitrogen and phosphorus content per mass (Nmass, Pmass), carbon discrimination rate (D13C), maximum rate of carboxylation per mass (Vcmax), time-integrated photosynthetic rate per mass (Amass), vein length per area (or vein density; VLA), stomatal density (d), stomatal initiation rate (i), and anatomical maximum stomatal conductance (gmax).

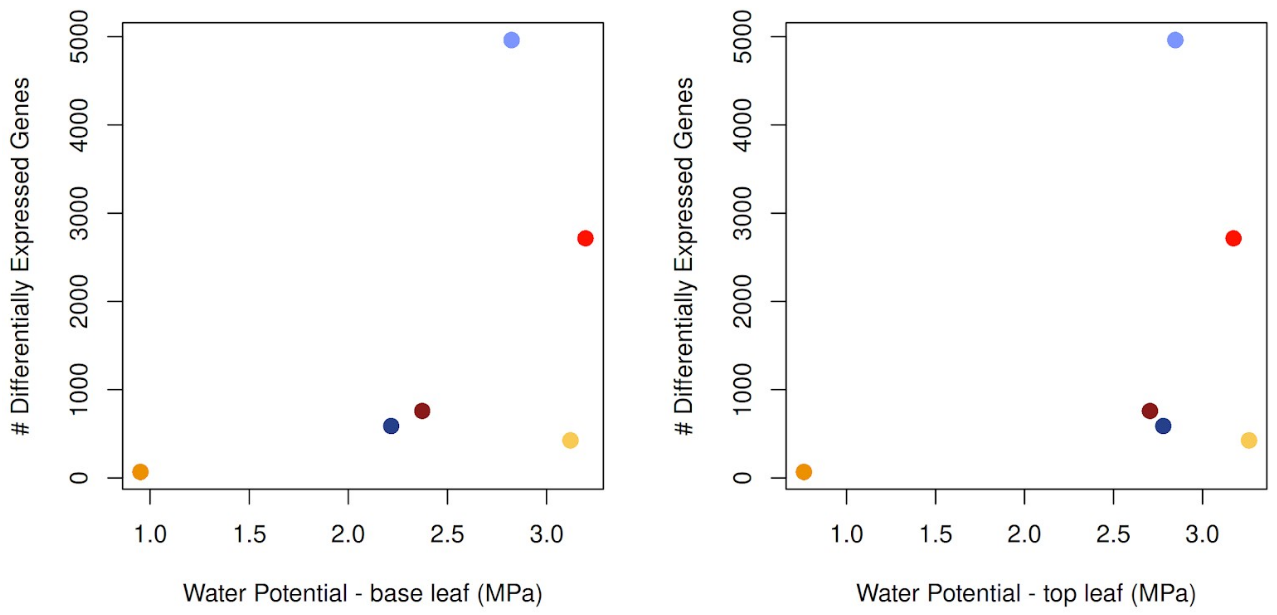


80 **Figure S6.** Species average leaf water potential of treated leaves and the differential expression
81 response. Two measurements were taken for each branch; one from a leaf at the base of the branch
82 (left), and one from the top leaf (right).

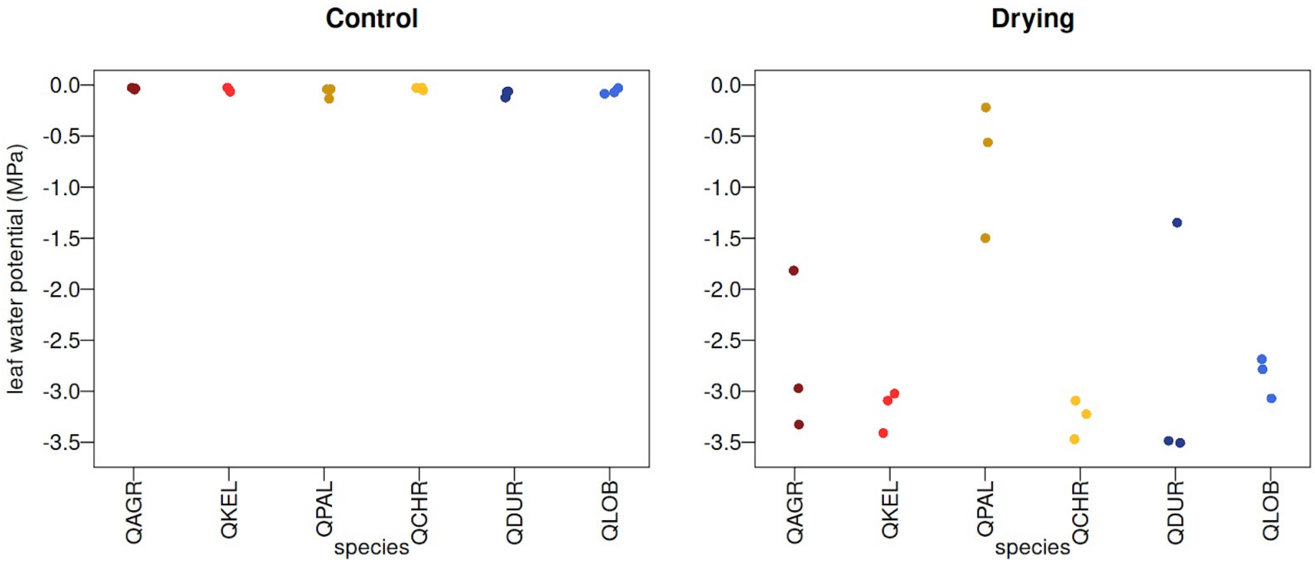


**Figure S7.** Leaf water potential for sequenced individuals in the control and drying treatment. Each point shows the measurement for one leaf from a branch. After the drying treatment, the leaf water potential of *Q. palmeri* was significantly different from other species (pairwise t-test, $p<0.05$); while all other pairs were not statistically different from each other.
